# A major QTL for resistance against Salmonid Rickettsial Septicaemia in coho salmon (*Oncorhynchus kisutch*) maps to a narrow region on chromosome 21, implicating two candidate genes

**DOI:** 10.64898/2025.12.08.689227

**Authors:** Thomas Moen, Fabian Grammes, Victor Martinez, Jacob Torgersen, Jørgen Ødegård, Tim Martin Knutsen, Tomasz Podgorniak, Robert Deerenberg, Daniela Cichero

## Abstract

The facultative intracellular bacterium *Piscirickettsia salmonis* causes Salmon Rickettsial Syndrome (SRS) in Coho salmon, Atlantic salmon, and other salmonids. SRS causes large mortalities in Chilean aquaculture and leads to heavy usage of antibiotics. In Atlantic salmon SRS resistance is a polygenic trait with moderate to large heritability. In coho salmon a large QTL for SRS-resistance was earlier found on chromosome 21. In the present study we have further characterized genetic resistance to SRS in coho salmon and searched for putative candidate genes underlying the QTL.

The mean heritability of survival was 0.31 and 0.58 on the observed and liability scale, respectively. The QTL on chromosome 21 explained from 26% to 97% of genetic variation within 12 different datasets. Two SNPs were substantially more significant compared to other SNPs and in very strong linkage disequilibrium with each other. The resistance allele was found to be dominant over the susceptibility allele at these SNPs.

One of the two SNPs was located within the first exon of two genes which are transcribed in opposite directions: a histidine triad nucleotide-binding protein 3 (*hint3*) gene and a gene (LOC109866666) encoding a long non-coding RNA (lncRNA). Genotypes at the SNP were correlated with expression levels at both *hint3* and the lncRNA gene, and the differential expression was manifested in both SRS-challenged and non-challenged fish. The exonic SNP is located 3 base pairs upstream of the start codon of the *hint3* gene, at a position which is crucial for effective translation according to the rules of Kozak. The resistance allele at the SNP correlates to increased expression levels and increased translation levels at *hint3*, although the latter remains to be experimentally proven.

Thus, it seems plausible that the QTL is due to the action of the *hint3* gene and/or the gene lncRNA gene encoded by LOC109866666. The hint3 gene on chromosome 21 is different from homologs in Atlantic salmon, and no Atlantic salmon homologs of LOC109866666 were found. Thus, it might be possible to increase SRS-resistance of Atlantic salmon by inserting the coho gene(s) through gene editing.

## Introduction

Piscirickettsiosis (SRS) is a bacterial disease caused by the gram-negative, facultative intracellular bacterium *Piscirickettsia salmonis.* The disease affects the farmed salmonids, coho salmon (*Oncorhynchus kisutch*), Atlantic salmon (*Salmo salar*) and rainbow trout (*Oncorhynchus mykiss*), but other salmonids can also be affected. While SRS outbreaks are reported from most salmonid farming countries (Norway, Canada, Scotland, Ireland, and Chile), the impact of SRS is especially large on coho salmon and Atlantic salmon in Chile. In 2023, 2.2% of mortalities in Chilean farmed coho were due to SRS, a decline from 7.2% in 2023. In the case of Atlantic salmon, 44.7% of mortalities were due to SRS in 2023 (Servicio Nacional de Pesca y Acuicultura 2024).

SRS typically appears in post smolts, two to three months after seawater transfer and is characterized by poor growth, atypical behaviour, darkened skin with haemorrhages, as well as gill lesions characterized by hyperplasia and fusion of the lamella (Almendras and Fuentealbas 1997). Internally, a pale and mottled liver is a typical sign, as is a swollen and discoloured kidney and enlarged spleen. Histopathological changes commonly include necrosis and inflammation in the liver, spleen, kidney, and intestine. Cardiac pathological signs include pericarditis, endocarditis, and focal myocardial necrosis. *Piscirickettsia salmonis* infects a broad range of host cells, particularly circulating macrophages, where it replicates within membrane-bound intracytoplasmic vacuoles of variable size. According to recent findings, the bacterium is not an obligate pathogen, but a bacterium with disease-causing potential that is often resident in the environment and/or the normal salmon microbiome without consistently producing disease (Cabello et al. 2019).

Since the first major Chilean SRS outbreak in 1989, various strategies have been implemented to prevent or control SRS. These strategies include constant disease monitoring, stress reduction, the use of immunostimulants, antibiotic treatments, vaccination and selective breeding. Several commercial vaccines are currently available against SRS in Chile, but their efficacy in the field is often low (Valenzuela-Avilés et al. 2022).

Selective breeding can be a good supplement or alternative to vaccination or other measures. Previous studies have demonstrated significant variation in resistance to *Piscirickettsia salmonis* in Atlantic salmon (*Salmo salar*), rainbow trout (*Oncorhynchus mykiss*) and coho salmon (*Oncorhynchus kisutch*) using disease challenge data, with heritability estimates ranging from 0.14 to 0.88 (Bangera et al. 2017, Barría et al. 2019, Moraleda et al. 2021, Yoshida et al. 2018, Yáñez et al. 2013, Yáñez et al. 2016, Yáñez et al. 2019). In Atlantic salmon and rainbow trout, resistance to SRS is a polygenic trait (Marín-Nahuelpi 2024, Barría et al. 2019), but in coho salmon a major QTL has been identified (Moen et al. 2022). Here, we present further results regarding this QTL and the genetic control of SRS-resistance in coho salmon.

## Results

Challenge tests were performed on six different year classes (yc) of coho salmon belonging to AquaGen: 2014, 2016, 2017, 2018, 2019, and 2021. Even year classes came from one breeding nucleus (population), odd year classes came from another. A cohabitant test model was used where mortality rates ranged from 0.67 to 1.00 in cohabitants and from 0.39 to 0.85 in shedders (Figure 1, Table 1).

**Figure 1:**
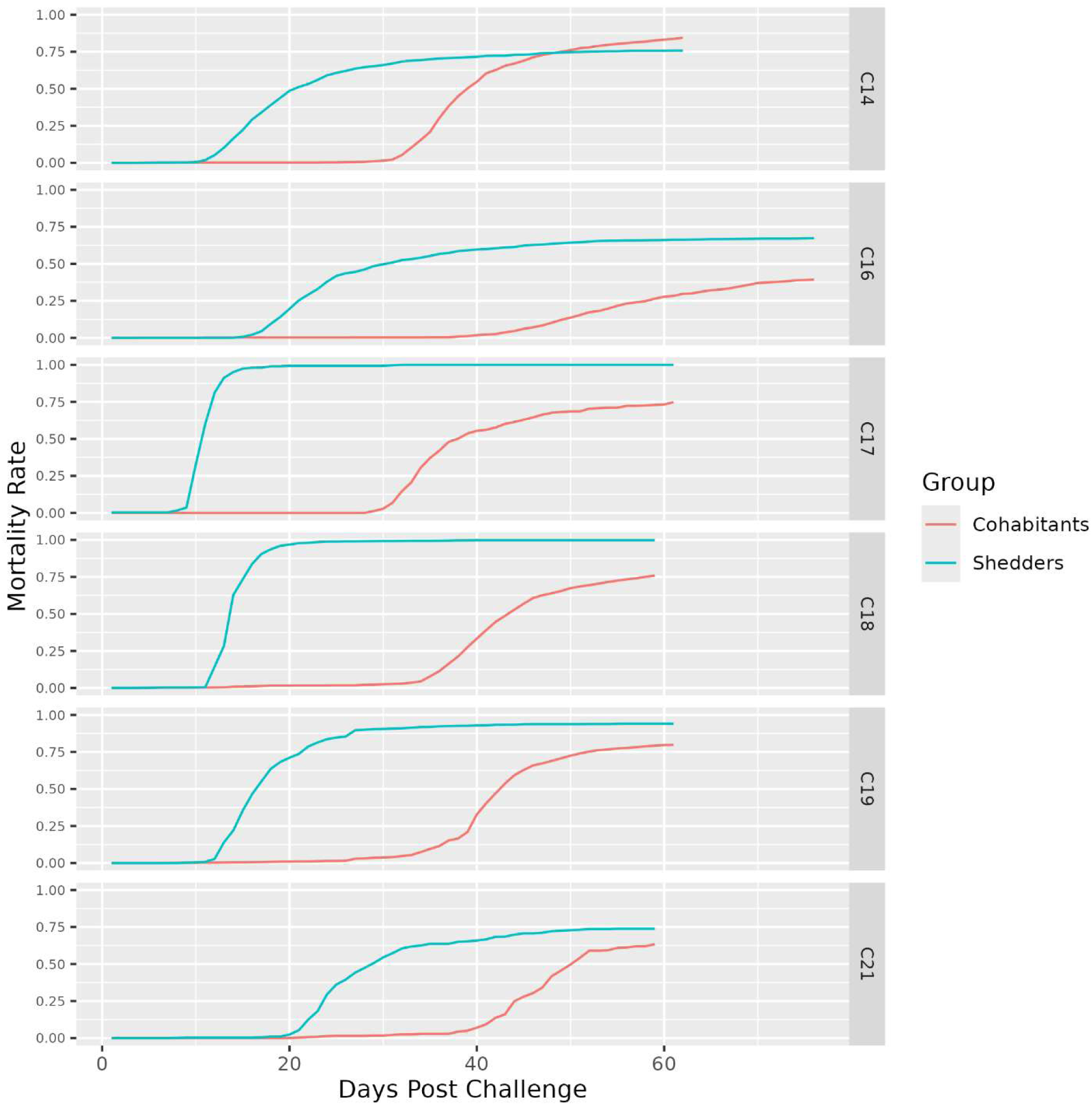
Mortality curves from challenge-tests.

**Table 1:** The datasets of the study. Each dataset is named according to year class (C14 for the 2014 year-class etc.) and whether the dataset consists of shedders (sh) or cohabitants (co). N = number of fish with phenotypes and genotypes (the aim was to genotype all challenge-tested fish, but some dropped out because of poor genotype quality etc). N mort., N surv. = Number of survivors and mortalities at the end of challenge test. h^2^ DPC, h^2^ survival observed, h^2^ survival liability = heritabilities for days-post-challenge, binary survival on the observed scale and binary survival on the underlying liability scale, all presented with standard errors *This dataset was not genotyped; the numbers are for challenge-tested fish. Body weights were measured at the end of the challenge test, for all datasets except C19co, C21co, and C21sh. For those three datasets, the listed body weights were measured prior to challenge.

| Data set | N | Weigh<br>ht (g) | N<br>Mort | N<br>Surv. | Mort.<br>Rate | h<sup>2</sup> DPC | h<sup>2</sup> Survival<br>Observed | h<sup>2</sup> Survival<br>Liability |
| --- | --- | --- | --- | --- | --- | --- | --- | --- |
| C14sh | 1062 | 299 | 805 | 257 | 0.76 | 0.44 ± 0.05 | 0.33 ± 0.05 | 0.62 ± 0.09 |
| C14co | 1011 | 305 | 855 | 156 | 0.85 | 0.49 ± 0.05 | 0.35 ± 0.05 | 0.81 ± 0.11 |
| C16sh | 1401 | 495 | 943 | 458 | 0.67 | 0.44 ± 0.04 | 0.36 ± 0.05 | 0.59 ± 0.08 |
| C16co | 1267 | 657 | 499 | 768 | 0.39 | 0.24 ± 0.04 | 0.25 ± 0.04 | 0.41 ± 0.07 |
| C17sh | 308 | 178 | 308 | 0 | 1.00 | 0.16 ± 0.08 | NA | NA |
| C17co | 321 | 267 | 240 | 81 | 0.75 | 0.55 ± 0.09 | 0.36 ± 0.10 | 0.68 ± 0.18 |
| C18sh | 1416 | 126 | 1415 | 1 | 1.00 | 0.26 ± 0.04 | NA | NA |
| C18co | 1392 | 219 | 1054 | 338 | 0.76 | 0.38 ± 0.04 | 0.38 ± 0.04 | 0.70 ± 0.08 |
| C19sh* | 1517 | 261 | 1436 | 81 | 0.95 | NA | NA | NA |
| C19co | 1350 | 246 | 1064 | 286 | 0.79 | 0.08 ± 0.03 | 0.08 ± 0.03 | 0.16 ± 0.06 |
| C21sh | 600 | 194 | 444 | 156 | 0.74 | 0.42 ± 0.07 | 0.30 ± 0.06 | 0.54 ± 0.11 |
| C21co | 600 | 196 | 383 | 217 | 0.64 | 0.41 ± 0.06 | 0.42 ± 0.07 | 0.70 ± 0.11 |

**Table 2:** Comparing performance of the overall best tag-SNP versus the best tag-SNP in individual dataset. Best-SNP from Meta-GWAS = The overall best tag-SNP, i.e. the most significant SNP from the GWAS (Affx-1264130923). Best-SNP from individual GWAS = The most significant SNP from GWAS in the individual dataset in question. SNP-effect = Allele substitution effect P-value = p-value from GWAS Frac. Genet. Var. = Fraction of genetic variance explained by the SNP in the dataset in question. Numbers are missing in the three leftmost columns when the overall best tag-SNP is the best tag-SNP in the dataset in question.

|  | Best-SNP from Meta-GWAS |  |  | Best-SNP from individual GWAS |  |  |
| --- | --- | --- | --- | --- | --- | --- |
|  | SNP-effect | p-value | Frac. Genet. Var. | SNP-effect | p-value | Frac. Genet. Var. |
| C14co | 11.4 | 3.3E-69 | 0.75 | - | - | - |
| C14sh | 13.5 | 1.7E-31 | 0.38 | - | - | - |
| C16co | 8.0 | 4.6E-22 | 0.37 | - | - | - |
| C16sh | 18.0 | 1.3E-27 | 0.26 | - | - | - |
| C17co | 15.9 | 3.0E-22 | 0.69 | 9.5 | 6.9E-24 | 0.45 |
| C18co | 9.1 | 4.2E-63 | 0.76 | - | - | - |
| C18sh | 2.3 | 2.3E-47 | 0.74 | - | - | - |
| C19co | 5.3 | 1.2E-11 | 0.51 | 5.3 | 2.6E-13 | 0.53 |
| C21co | 10.3 | 7.7E-31 | 0.97 | 10.4 | 6.4E-32 | 1.00 |
| C21sh | 16.5 | 1.3E-22 | 0.64 | 16.1 | 7.4E-23 | 0.61 |

In shedders, the mean heritabilities (across datasets) for days-post-challenge (DPC), survival on the observed scale, and survival on the liability scale were 0.34, 0.33, and 0.58, respectively. For cohabitants the same means were 0.36, 0.31, and 0.58 (Table 1). Among the cohabitant datasets, the dataset from yc 2019 was an outlier in terms of having low heritability. Expecting low heritability also in shedders from yc 2019, we decided not to genotype shedders from that year-class. In the 2017 year-class, heritability was high among cohabitants but not statistically different from zero among shedders. This was probably due to a very high infection pressure (i.e. steep mortality curve) among shedders in that year-class (Figure 1). Consequently, shedders from yc 2017 were not used in subsequent analyses.

Genetic correlations between DPC in cohabitants and DPC in shedders were high: 0.84 ± 0.06 for yc 2014, 0.96 ± 0.06 for yc 2016, 0.87 ± 0.07 for yc 2019, and 1.00 ± 0.09 for yc 2021. Except for yc 2014, these estimates are not significantly different from 1. Thus, data from shedders and cohabitants can be used on equal footings to make inferences about the genetics of SRS resistance in coho (at least in these datasets).

Estimates for the genetic correlation between DPC and binary survival were calculated in datasets C14co, C14sh, C16co, C16sh, C17co, C18co, C21co, and C21sh (co = cohabitants, sh = shedders). The estimates (± standard error) were 0.96 ± 0.02, 0.98 ± 0.01, 1.00 ± 0.01, 1.00 ± 0.0, 0.97 ± 0.03, 0.99 ± 0.01, 1.00 ± 0.02, and 1.00 ± 0.01, respectively. None of the estimates were significantly different from 1. It is therefore reasonable to conclude that DPC and survival are reflective of the same biological trait in these datasets.

A meta-GWAS performed on the entire dataset was dominated by a large QTL on chromosome 21. The QTL disappeared when the best tag-SNP for the QTL was included as cofactor in the model, confirming that the tag-SNP captures the QTL very well (Figure 2). Zooming in on chromosome 21, it became clear that the strongest GWAS signals were localised to a relatively narrow genomic region (Figure 3). The region contained two SNPs which were much more significant than other SNPs: Affx-1264130923 (position in NC_034194.2 = 22,837,421, p-value = 4.7e-264, allele substitution effect = 9.96) and Affx-1264130925 (position in NC_034194.2 = 22,842,793; p-value = 2.4e-263, allele substitution effect = 9.95). The two SNPs were in almost perfect linkage disequilibrium (LD) with each other, and the few deviations that were found between them may have been due to genotyping errors.

**Figure 2:**
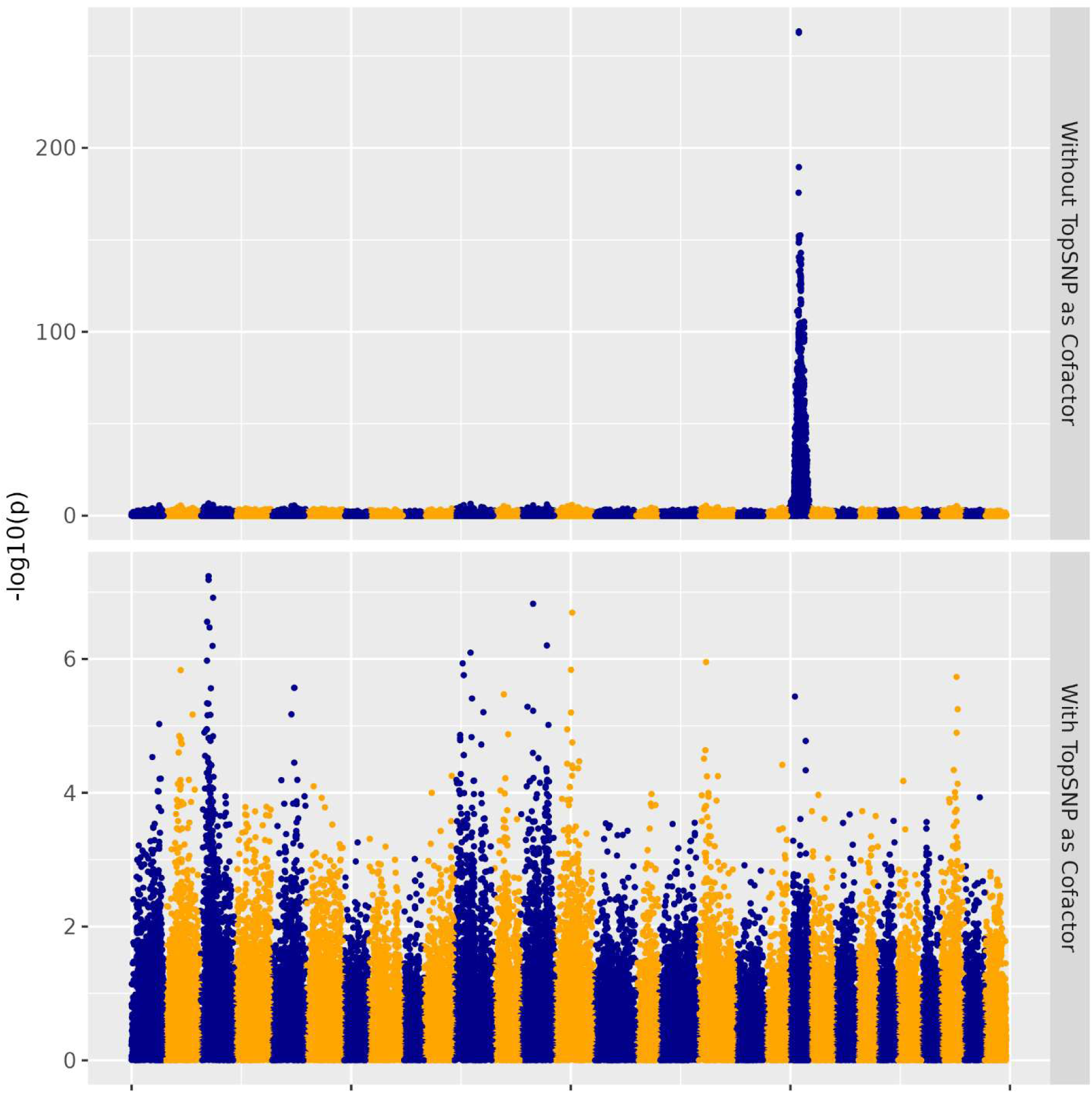
Meta-GWAS. Upper panel: The genotype of Affx-1264130923 was not included as cofactor in the mode Lower panel: The genotype of Affx-1264130923 was included as cofactor in the mode

**Figure 3:**
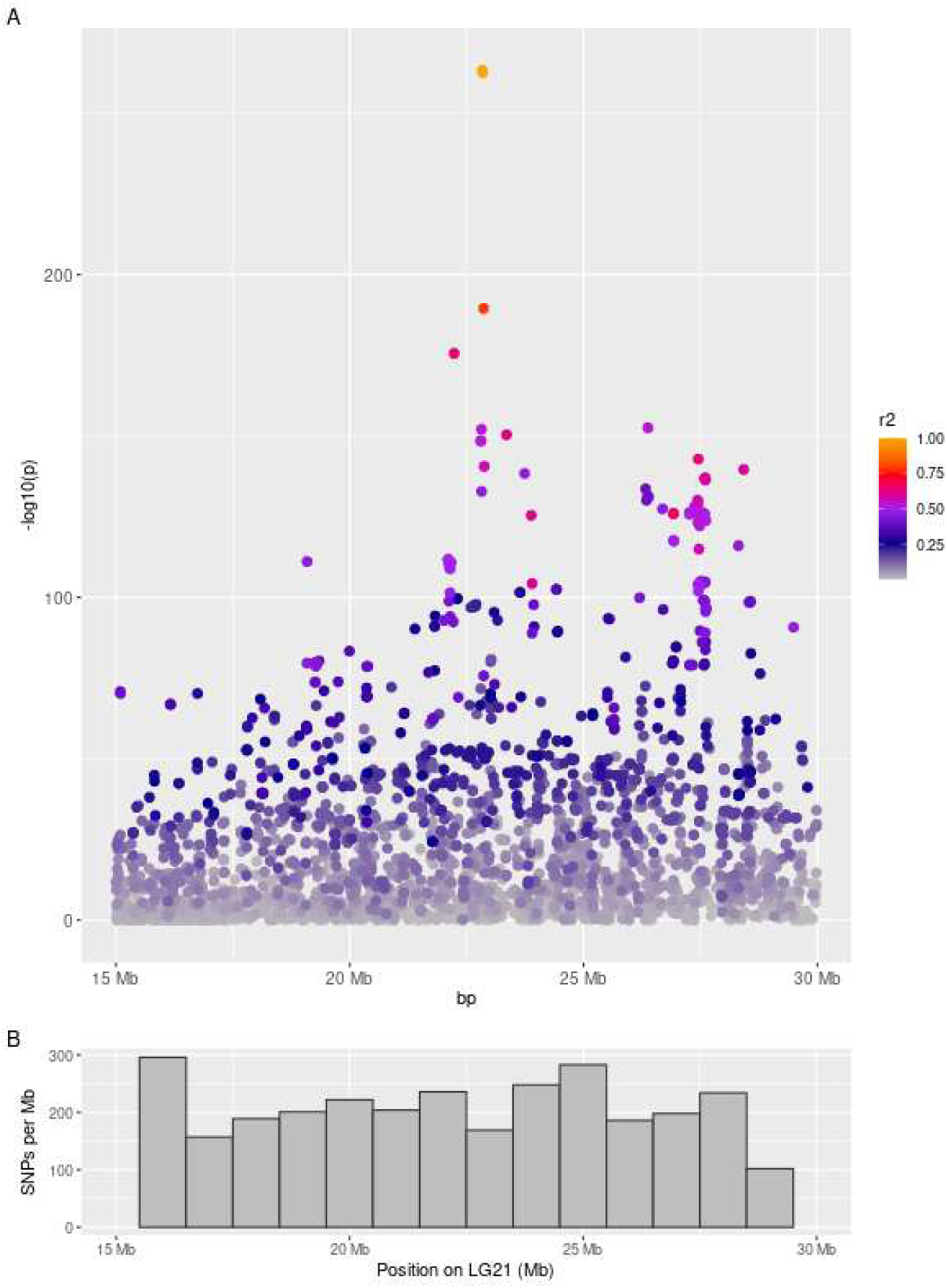
GWAS, zooming in on chromosome 21. The bottom panel displays the SNP-density in the data

GWASes run on individual datasets showed that Affx-1264130923 and/or Affx-1264130925 was the best tag-SNP in five out of nine individual datasets. In the remaining four datasets, Affx-1264130923/Affx-1264130925 was almost as significant as the most significant SNP within that dataset. The deviations were arguably so small they could be caused by randomness in the data. Notably, the deviations occur mainly in datasets which have relatively small sample sizes (C17co, C21co, C21sh) or low heritability (C19co). The amount of randomness would be largest in these datasets. It should also be noted that Affx-1264130923/Affx-1264130925 were shown to be the best tag-SNPs in two coho populations unrelated to the AquaGen breeding nuclei (data not shown).

The ability of Affx-1264130923 and Affx-1264130925 to predict the trait is illustrated in Figure 4, which displays the relationship between survival rate and the number of resistance alleles at Affx-1264130923. The figure indicates a substantial dominance effect at the resistance allele (Q). One dataset indicates overdominance, but it must be noted that few fish are homozygous for the resistance allele, due to low allele frequency. Thus, the observed overdominance is most likely an artefact of few datapoints within the QQ group.

**Figure 4:**
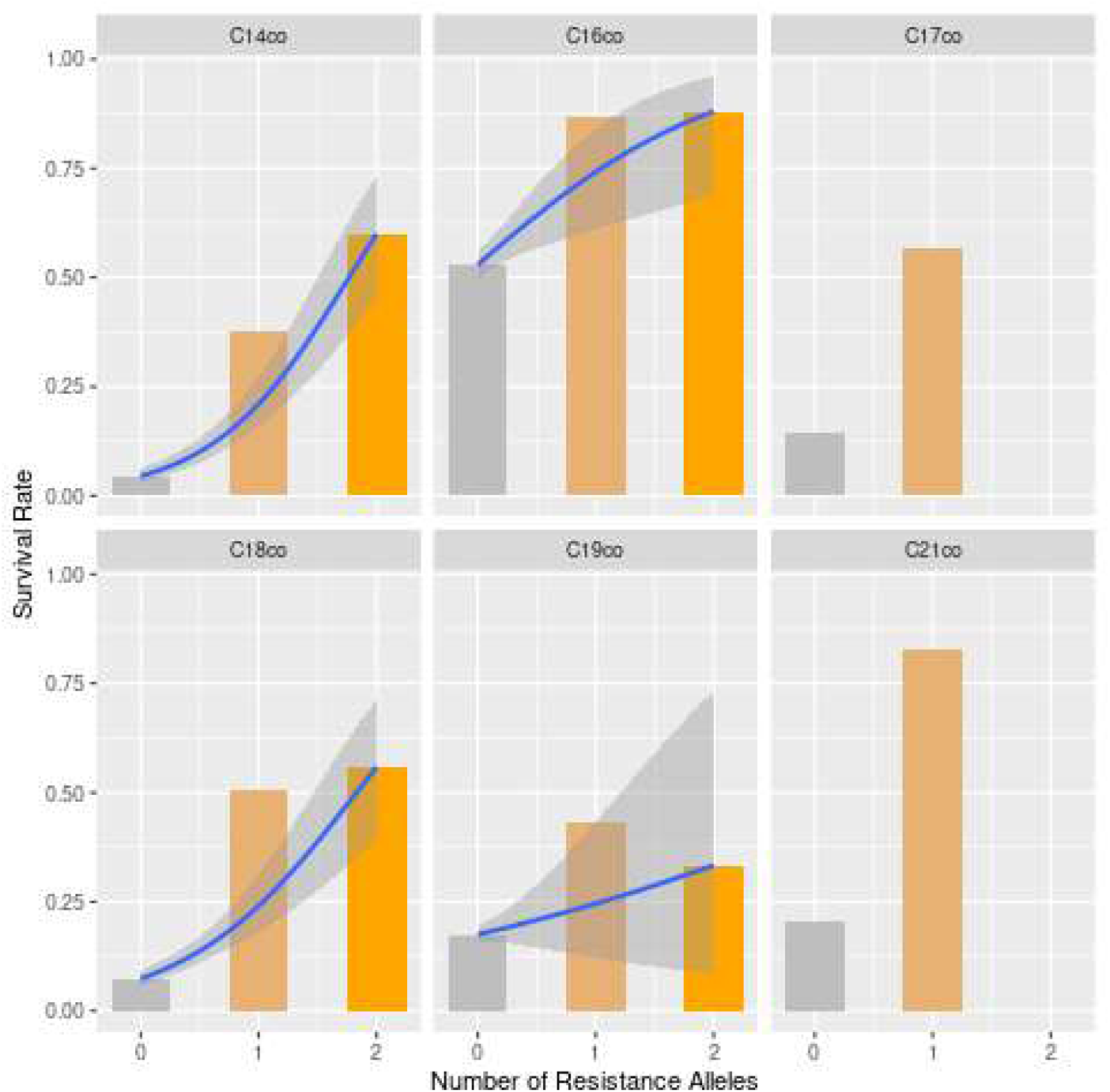
Relationship between survival rate and number of resistance alleles (Q) at the QTL. The tag-SNP used is Affx-1264130923, the most significant SNP. The line is a logistic regression line formed by the data points at the homozygous genotypes, and the shaded area is the 95 % confidence interval of that line. The positioning of the heterozygous genotypes outside the 95 % confidence interval indicates dominance at the QTL.

It is not surprising that Affx-1264130923 and Affx-1264130925 are very strong tag-SNPs for the QTL, because they were the end-product of a fine-mapping exercise in which whole-genome sequencing (WGS) was used to identify candidate loci, whereupon all loci of interest were tested out on an updated SNP-chip (Moen et al. 2022). However, in that fine-mapping process, SNPs of interest may have been overlooked for several reasons: 1) The genotype contrast used for finding candidate loci in the WGS data was only an approximation of the unknown causative mutation (Quantitative Trait Nucleotide, QTN). 2) Some SNPs of interest might not have been genotyped on the SNP-chip because their flanking DNA sequence did not permit SNP-chip genotyping or because the genotype quality from the SNP-chip was too low. Therefore, SNPs of interest might still be hidden in the WGS data. To find out, we searched the WGS data for SNPs in medium to strong LD with Affx-1264130923. The results showed that all SNPs in very strong LD with Affx-1264130923 were already on the SNP-chip, and that LD (r^2^) values depended strongly on physical distance to Affx-1264130923 (Figure 5). Thus, the WGS data did not reveal additional SNPs which could rival Affx-1264130923 or Affx-1264130925 as tag-SNP or as QTN candidates.

**Figure 5:**
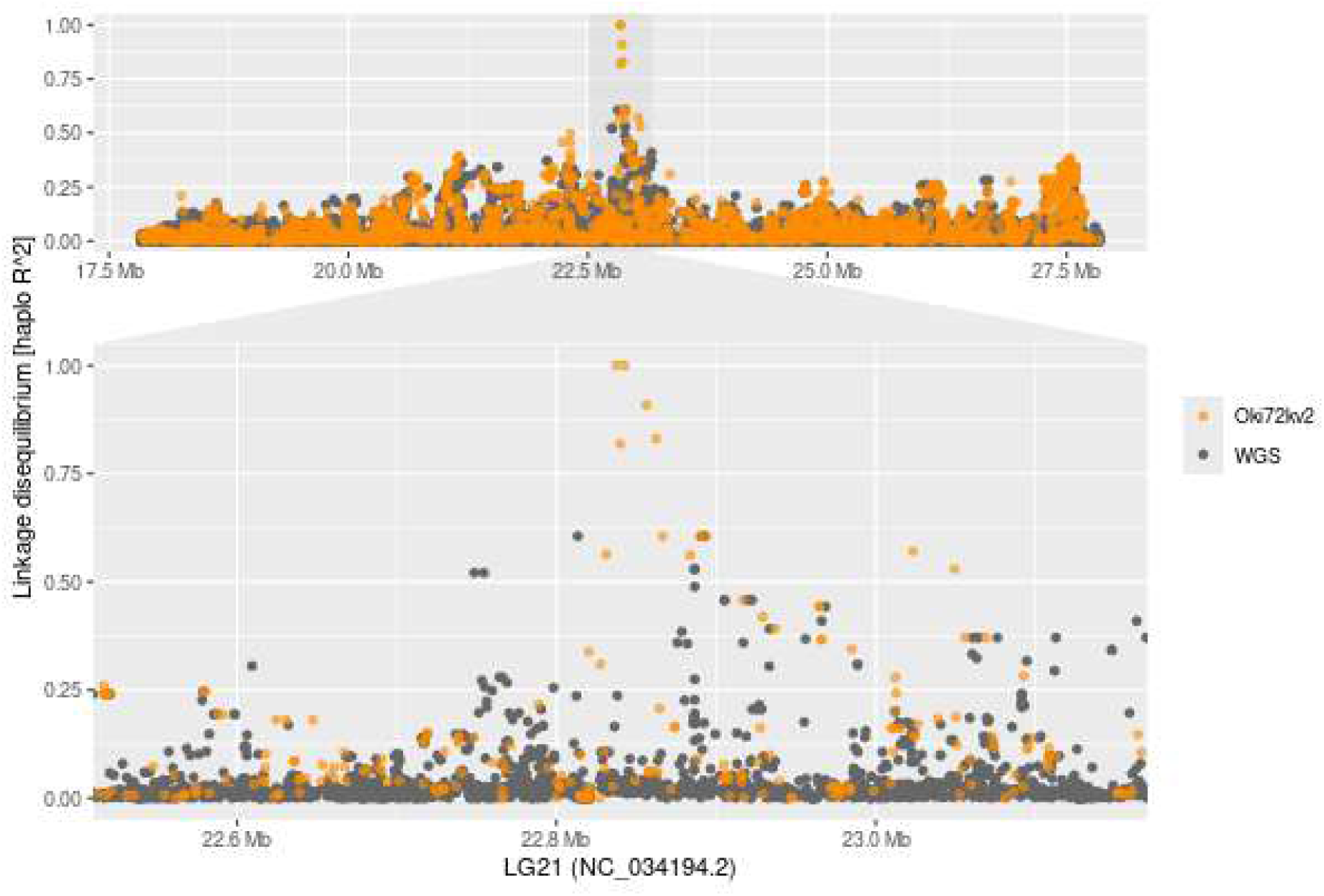
GWAS: LD between the best tag-SNP (Affx-1264130923) and WGS-derived SNPs. The figure is based on WGS on a diverse sample of 91 fish from the AquaGen breeding nuclei

Given the apparent high precision of the estimate for QTL position, it is reasonable to look to the genomic region immediately surrounding Affx-1264130923/Affx-1264130925 to identify candidate causal genes. Figure 6 zooms in on the QTL and overlays the genes found in the region according to the RefSeq gene annotation. Affx-1264130923 is located in the first intron of a gene annotated as *Histidine triad nucleotide-binding protein 3* (*hint3, LOC109866665*), 22 bp from the start of the second exon. Affx-1264130925 is located within the 5’ UTR of *hint3*, but also within the first exon of an overlapping gene, with identifier LOC109866666 (Figure 7). That locus encodes a long non-coding RNA (lncRNA), and the first exon stretches from position 22,842,549 to position 22,842,859. Downstream of LOC109866666 lies a gene annotated as *Nuclear receptor coactivator 7* (*ncoa7,* LOC109866653). An investigation of the expression of these three genes in a tissue panel showed that the lncRNA is the most widely and highest expressed among the three genes. *Hint3* showed medium expression in brain, gill, and gonad, but low expression in head kidney and spleen. *Ncoa7* displayed high expression in liver and low expression in head kidney, gonad, and distal intestine (Figure 8). We noticed that gene models differed between RefSeq and Ensembl: While *ncoa7* shows good correspondence, Ensembl is lacking the lncRNA according to Ensembl. *Hint3* is truncated and lacks several exons as well as the 3-UTR in the Ensemble annotation. For the three genes within the narrow QTL region the RefSeq gene models showed better correspondence the RNAseq read alignments.

**Figure 6:**
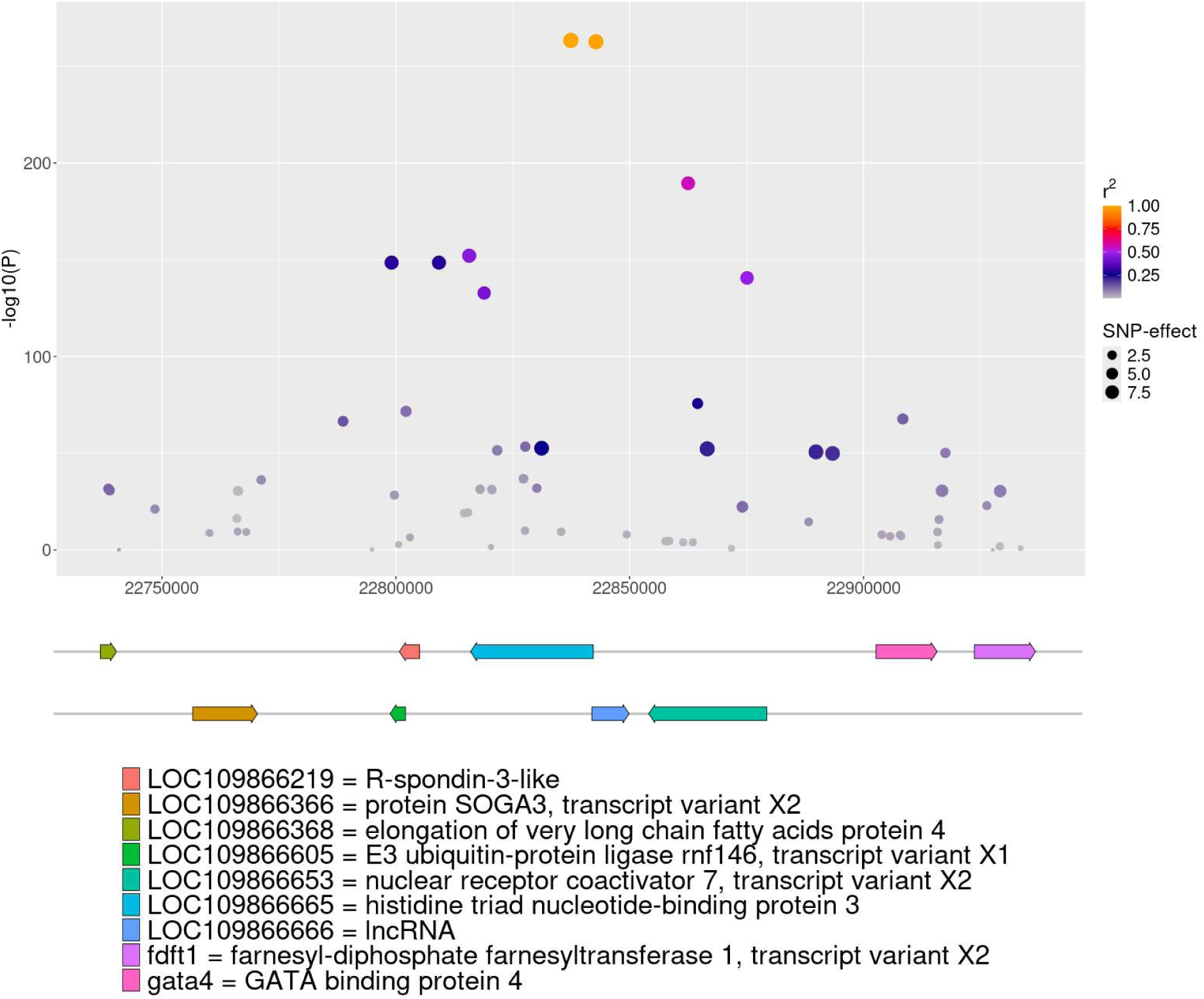
Candidate genes in the immediate QTL region. Gene names are according to the annotations of coho genome reference GCF_002021735.2 (NCBI annotation release 101 and annotation release 114 from EMBL-EBI (via Ensembl)). In the GWAS plot, SNPs are coloured according to LD (r^2^) to Affx-1264130923; r^2^ was calculated across a combined dataset encompassing all year classes of Table 1. The size of SNPs in the GWAS plot are according to the magnitude of the allele substitution effect.

**Figure 7:**
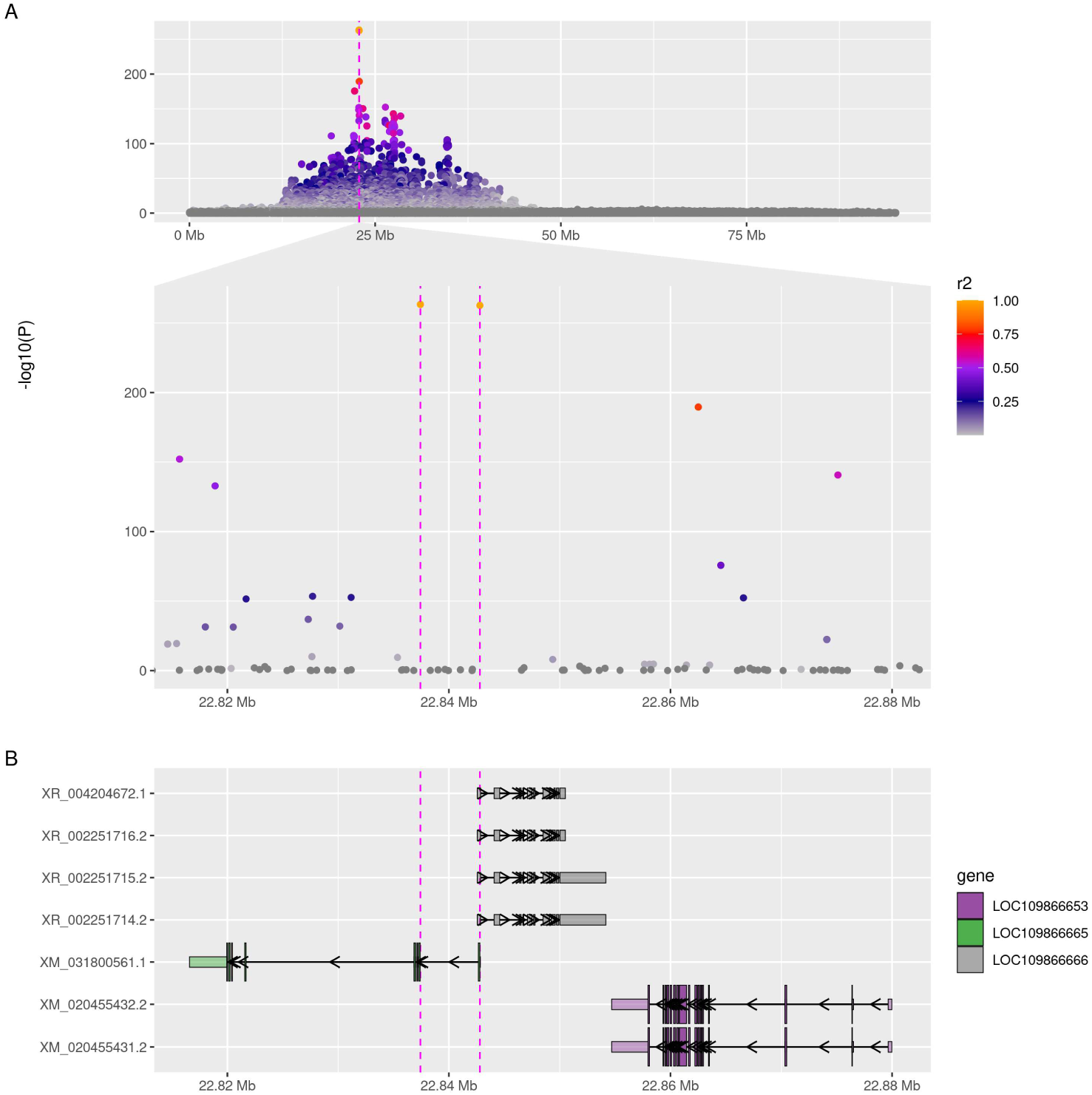
Genomic evidence for the SRS QTL region on LG21. **A**: Manhattan plot for the meta-GWAS of SRS resistance with a magnification to the genomic region (LG21: 22.816.584-22.880.322) containing the two top SNPs. **B**: RefSeq (<u>GCF_002021735.2</u>) based gene annotations.

**Figure 8:**
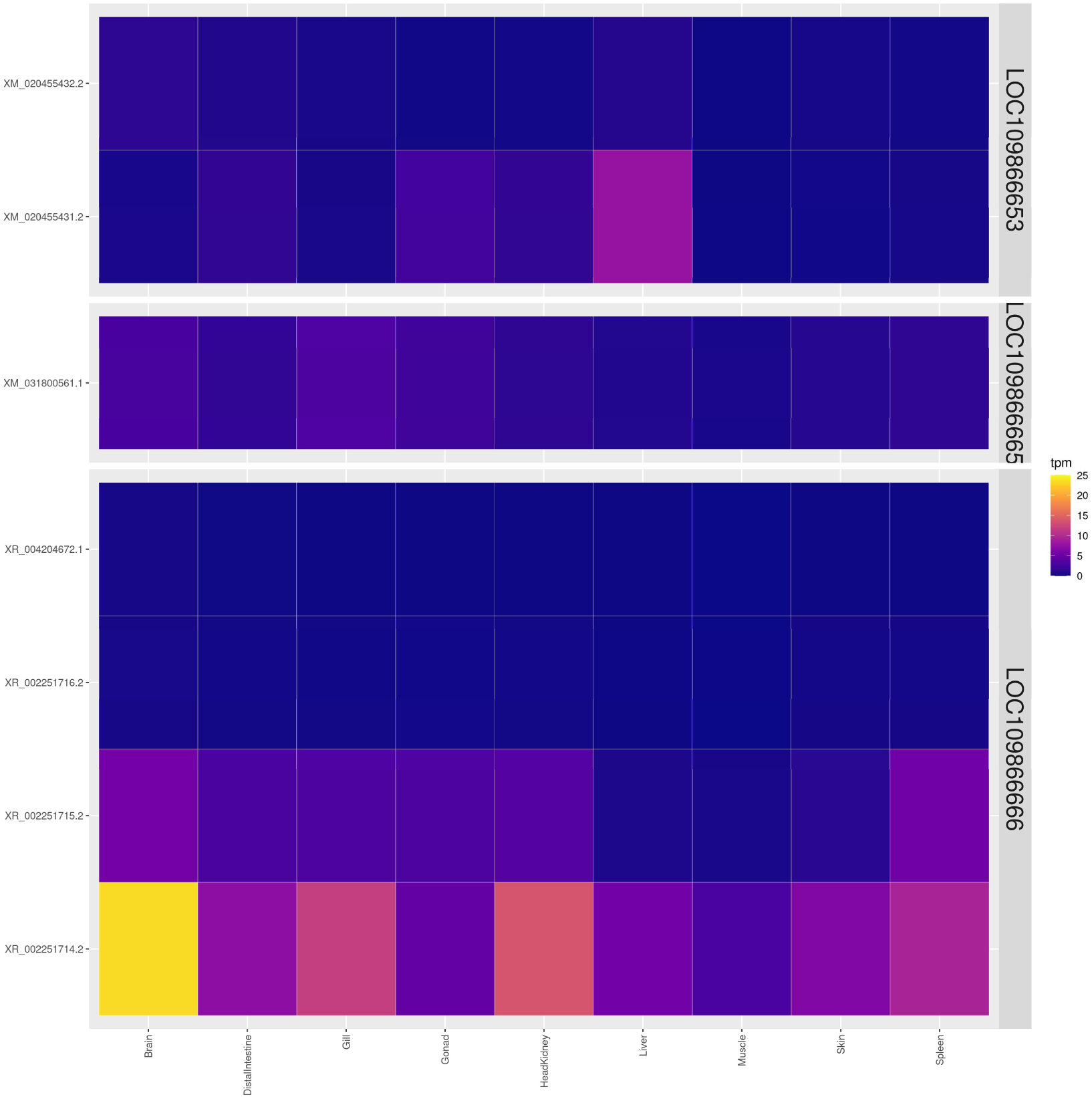
Heatmap showing the expression levels for all transcripts of the 3 genes in close proximity to the QTL. *hint3:* LOC109866665 *lncRNA:* LOC109866666 n*coa7:* LOC109866653

A cis-eQTL analysis was performed to investigate the positional candidate genes further. The cis-eQTL analysis based on the yc 2014 challenge trial identified a single significant association after stringent FDR correction (α = 0.05): Affx-1264130925 exhibited the strongest regulatory signal.

Genotypes at the SNP were significantly correlated with expression levels at *hint3* (LOC109866665; p-value = 0.003; FDR-adjusted p-value = 0.02; Table 3). The SNP also appeared correlated with expression levels at the lncRNA (LOC109866666) and the gene *soga3* (LOC109866366), although these correlations were not statistically significant. Genotypic model comparison revealed that a dominant genetic model (contrary to expectation) provided the best fit, supported by both the lowest Akaike Information Criterion (AIC) and highest statistical significance among additive and dominant models (Table 4).In the best-fitting dominant model, the presence of at least one alternative allele was associated with a 1.80-fold increase in *hint3* expression (95% CI: 1.26–2.58) compared to reference homozygotes.

**Table 3:** Cis-eQTl analysis of genes spanning 100 kb upstream and downstream from Affx-1264130925. FDR = False Discovery Rate

|  |  | Additive |  |  | Dominance |  |  |
| --- | --- | --- | --- | --- | --- | --- | --- |
| Gene ID | Distance<br>in bp | Fold<br>change<br>per allele | P-value | FDR-adjusted<br>p value | Fold<br>change<br>per allele | P-value | FDR-adjusted<br>p value |
| LOC109866665 | 26209 | 1.49 | 0.01 | 0.05 | 1.8 | 0.0032 | 0.0192 |
| LOC109866666 | 244 | 1.68 | 0.04 | 0.09 | 1.5673 | 0.0655 | 0.1324 |
| LOC109866366 | 85607 | 1.73 | 0.05 | 0.09 | 1.7815 | 0.0662 | 0.1324 |
| LOC109866605 | 43418 | 1.30 | 0.19 | 0.27 | 1.3057 | 0.1491 | 0.2236 |
| LOC109866653 | 11898 | 1.17 | 0.45 | 0.55 | 1.1708 | 0.4791 | 0.5749 |
| fdft1 | 81541 | 0.94 | 0.85 | 0.84 | 0.8553 | 0.5973 | 0.5973 |

**Table 4:** Nuclotide BLAST hits with the cDNA sequence of LOC109866666 as query.

| Species | Scientific Name | Max Score | Total Score | Query Cover | E value | Per. ident | Acc. Len | Identifier of Hit |
| --- | --- | --- | --- | --- | --- | --- | --- | --- |
| Sockeye salmon | *Oncorhynchus nerka* | 3268 | 4497 | 0.86 | 0.0 | 96.29 | 3890 | XR_003865096.2 |
| Pink salmon | *Oncorhynchus gorbuscha* | 2602 | 4171 | 0.81 | 0.0 | 95.59 | 2695 | XR_006831875.1 |
| Chum salmon | *Oncorhynchus keta* | 2156 | 3395 | 0.65 | 0.0 | 95.75 | 5446 | XR_008140814.1 |
| Chinook salmon | *Oncorhynchus tshawytscha* | 1573 | 3426 | 0.63 | 0.0 | 98.35 | 2254 | XR_002948494.2 |
| Brown trout | *Salmo trutta* | 1412 | 3140 | 0.71 | 0.0 | 87.76 | 3670 | XR_003877641.1 |
| Arctic char | *Salvelinus alpinus* | 554 | 1185 | 0.24 | 9e-152 | 95.40 | 3433 | XR_011676432.1 |
| Masu salmon | *Oncorhynchus masou* | 348 | 348 | 0.12 | 7e-90 | 80.43 | 1738 | XR_010451722.1 |

Next, the eQTL analysis was extended to the RNAseq tissue panel, where expression of the different transcripts was modelled according to the genotype of the SNP Affx-1264130925. A significant effect of the SNP on the expression of *hint3* was found in gill, head-kidney and spleen and for one transcript of the lncRNA (XR_002251714.2) in gill (Figure 9).

**Figure 9:**
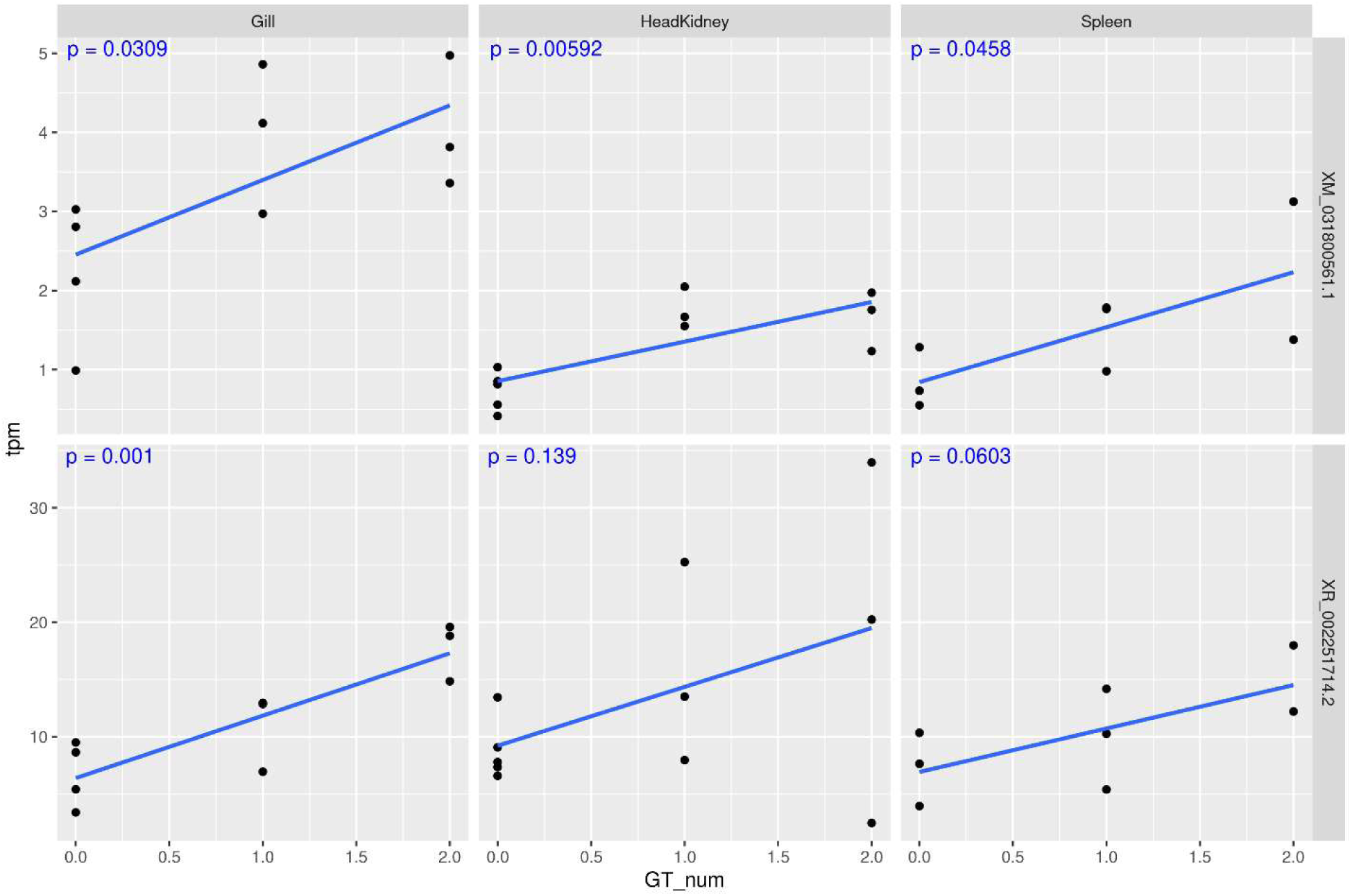
Tissue panel eQTL analysis: Effect of the SNP Affx-1264130925. SNP genotypes were derived from RNAseq on the transcript expression of the genes in the narrow QTL region. The eQTL p-value is displayed in the plot.

Although based on relatively small sample sizes, both eQTL experiments revealed a consistent effect of SNP Affx-1264130925 on expression of *hint3*. In all cases, the presence of the resistance allele was associated with increased expression of *hint3*. Similarly, both studies displayed a consistent effect of SNP Affx-1264130925 on expression of the lncRNA (LOC109866666), although that effect was not statistically significant in one of the studies (Table 3). These results support the notion that Affx-1264130925 functions as a cis-regulatory variant modulating transcription of both the lncRNA (LOC109866666) and *hint3* (LOC109866665) in head kidney, as well as spleen and gill both during the cohabitation challenge and under normal conditions. Most other neighbouring genes (e.g., gata4 and LOC109866219) exhibited negligible expression across samples and were therefore excluded from cis-eQTL consideration.

Affx-1264130925 has another highly likely impact on the *hint3* gene: The SNP lies 3 bp upstream of the start codon of the transcript, i.e. within the Kozak sequence surrounding the start codon. According to the Kozak consensus, an adenine (A) or a guanine (G) at the 3 bp upstream (-3 position) position is strongly preferred for effective translation, and as is a guanine at the +4 nucletide after the ATG (+4 position) (Kozak 1986). Because the resistance allele at Affx-1264130925 has an A at position -3 and the susceptibility allele has a T (relative to the read direction of the gene), transcripts carrying the resistance allele will be strongly translated whereas transcripts carrying the susceptibility allele would be weakly translated. Thus, the differential transcription at the *hint3* gene will be further amplified through translational differences.

## Discussion

In an earlier conference proceeding, we presented the identification of a major QTL for resistance to SRS in coho salmon. We also presented how novel SNPs were identified which could predict SRS-resistance across populations. In the present publication, we have investigated the precision of that fine-mapping, to understand whether our fine-mapping attempts managed to position the QTL at high precision, to understand the full potential of genomics-informed breeding for SRS-resistance in coho, and to identify putative candidate genes.

The results confirm that SRS-resistance in coho is a highly heritable trait, when measured using a controlled challenge test. Both shedders and cohabitants were informative in genomic predictions, although cohabitants were generally more informative. The genetic correlation between days-post-challenge and binary survival was not significantly different from 1, meaning that survivors should not be regarded as truly SRS-resistant fish; rather, they were tolerant enough to survive until the end of the test.

From 26% to 97% of the genetic variation in SRS-resistance was explained by the single major QTL on chromosome 21, across datasets. The resistance allele at the QTL had a strong dominance effect, meaning that much genetic progress can be achieved by only selecting on male selection candidates.

Allowing for a slight amount of randomness in the GWAS results, the two most significant SNPs were as powerful predictors of SRS-resistance as any other SNP on the SNP-chip in each of all 12 available datasets. The 12 datasets represented 4 distinct source populations of coho salmon, as the GWAS was performed on domesticated coho populations which were quite recently formed, based on four pre-existing domesticated populations. The findings were also confirmed in GWASes done on another two coho populations, unrelated to the ones used in the study (results not shown). Analysis of WGS data confirmed that no other SNPs in the genome were in strong LD with these most significant SNPs without themselves being part of the GWAS. Furthermore, a GWAS-approach based on haplotypes (across multiple SNPs) within the QTL region did not increase the fraction of genetic variance explained by the QTL (results not shown); haplotype-based approaches often help to ‘tag’ a QTL more precisely when available SNPs are not in strong LD with the QTN. Thus, there are strong reasons to believe that the identified SNPs are unique in their ability to predict SRS resistance in coho salmon. By extension, there are good reasons to believe that these SNPs are in physical proximity to the unknown QTN, or that one of them is the actual QTN.

The two most significant SNPs were located close to three genes: *hint3*, a lncRNA gene, and *ncoa7*. One of the SNPs, Affx-1264130925, is located within the 3’UTR of *hint3* and within the first exon of the lncRNA gene. Genotypes at the SNP was correlated to transcription levels of both *hint3* and the lncRNA, but not to *ncoa7.* The third most significant SNP caused an amino-acid change in Ncoa7, but that SNP was substantially less significant than the two top-SNPs. Consequently, *ncoa7* can probably be disregarded as a candidate gene for SRS resistance.

Affx-1264130925 was found to alter the Kozak sequence of the *hint3* gene, so that more Hint3 protein will be produced from *hint3* mRNAs carrying the resistance allele, compared to the susceptibility allele. The functional importance of the Kozak sequence is supported by extensive experimental evidence showing that variation in the Kozak context alters translational efficiency in human cells and other eukaryotic model organisms (e.g. Kozak 1986; Kozak 1987; Grzegorski 2014; Ambrosini 2022). A variety of single-gene human disorders are caused by haploinsufficiency, a genetic condition by which mutational inactivation of one allele leads to reduced protein levels and functional impairment. Such disorders include susceptibility to cancer, growth retardation, and developmental, neurological, or metabolic syndromes (e.g. Ambrosini 2022). Affx-1264130925 is expected to lead to a substantial difference in translational efficiency because the SNP is located at the most important position (-3) within the Kozak consensus. Thus, it seems likely that Affx-1264130925 can have a large impact on SRS-resistance if *hint3* is the causative gene. This pinpoints Affx-1264130925 as a possible candidate for being the QTN underlying SRS-resistance. Affx-1264130925 may have other functional implications in addition to the impact on the Kozak sequence. For example, we found that the SNP overlaps with a region of open chromatin according to ATAC-seq data (results not shown). Further research is needed to fully understand the functional impact of the SNP.

Hint3 belongs to the HIT (histidine triad) superfamily of protein and possible links to *Piscirickettsia Salmonis* resistance in coho include control of apoptosis or nutrient scavenging. In Rainbow trout macrophages *Piscirickettsia salmonis* induce apoptosis (Rojas et al. 2010) and altered Hint3 activity may affect cell death, by either preventing or promoting it, depending on the cellular context as shown in humans (Kang et al. 2025). The main function of Hint3 is production of AMP through hydrolysis of adenylated intermediates (Dolot et al. 2025). A possible function for Hint3 underlying *P. salmonis* resistance in coho salmon may be to limit the available AMP pool needed for bacterial replication. Rickettsiales lack the genetic machinery for *de novo* synthesis of nucleotides and as such rely on their hosts as a source. In addition to ADP and ATP, Rickettsiales also actively take up AMP which can be converted to ADP and ATP through phosphorylation (Atkinson et al. 1985).

There was also a tendency for Affx-1264130925 to be correlated with expression levels at the lncRNA. Long non-coding RNAs are RNAs longer than 200 bp which do not encode proteins. They have been found to have regulatory function, acting through mechanisms such as translational inhibition, mRNA degradation, recruitment of chromatin modifiers, regulation of protein activity, and by acting as RNA decoys. Most lncRNAs evolve more rapidly than protein-coding sequences, are cell type specific and regulate many aspects of cell differentiation and development and other physiological processes (reviewed in Mattick et al. 2023). LncRNAs also impact on immunity (reviewed in Bocchetti et al. 2021), and targeted delivery of lncRNA is being examined as a therapeutic against cancer (e.g. Coan et al. 2024). There are numerous examples of lncRNAs impacting on response to pathogens. For example, increasing evidence indicates that host lncRNAs play important roles in the interactions between mammalian hosts and *Mycobacterium tuberculosis*, the intracellular bacterium causing tuberculosis (reviewed in Cheng et al. 2025). As further examples, Boltaña (2016) et al. found widespread differential expression of lncRNAs in response to infection by Infectious Salmon Anemia (ISA) In Atlantic salmon, and Martinez et al. (2023) found differential expression at many lncRNAs during infection with *Piscirickettsia salmonis* in Atlantic salmon. Interestingly, the cDNA sequence corresponding to the lncRNA within the QTL region (identifier: LOC109866666) produced strongly matching nucleotide BLAST hits from a variety of salmonid species: Arctic char, brown trout, chinook salmon, chum salmon, masu salmon, pink salmon, and sockeye salmon (Table 4). In Atlantic salmon and rainbow trout, on the other hand, the gene was not found. A comparison of the coho salmon and Atlantic salmon genome references confirms the absence of this lcnRNA in Atlantic salmon (Figure 10).

**Figure 10:**
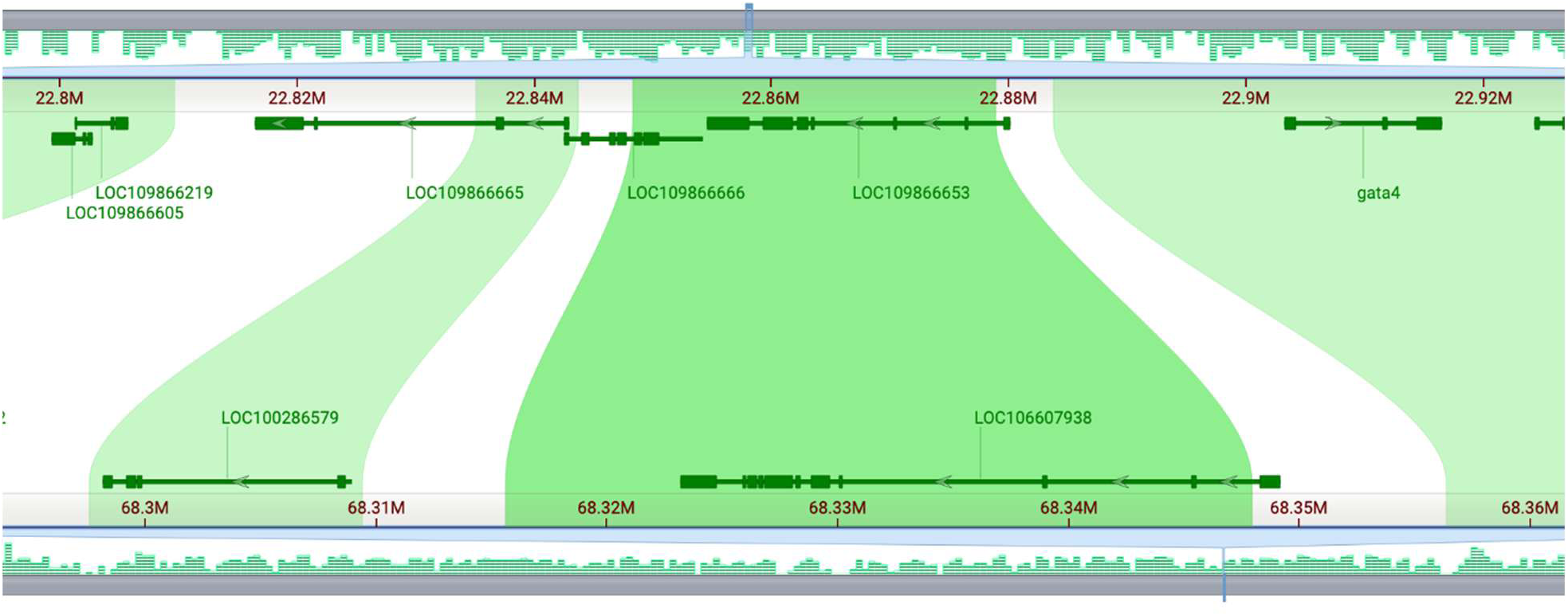
Comparative genomic alignment between Coho salmon and Atlantic salmon. Synteny analysis of the SRS-resistance QTL region generated using the NCBI Comparative Genome Viewer (Rangwala et al., 2024). The top track represents the Coho salmon (*Oncorhynchus kisutch*) genome (chromosome 21), and the bottom track represents the orthologous region in the Atlantic salmon (Salmo salar) genome. Green bands indicate regions of sequence homology. The alignment highlights that the candidate long non-coding RNA gene (LOC109866666), located between *hint3* (LOC109866665) and *ncoa7* (LOC109866653) in Coho salmon, is absent in the Atlantic salmon genome, while the flanking protein-coding genes show conserved synteny.

Our results indicate hint3 and/or the lncRNA encoded by LOC109866666 as putative causative genes underlying the QTL for resistance to SRS in coho salmon. In terms of selective breeding, identification of the causative gene(s) is not crucial, given that the already identified SNPs seem to capture the underlying causative mutation as accurately as possible. The latter statement, however, comes with a minor caveat, as the methodologies we have used have permitted the identification and genotyping of only SNPs and short insertion/deletions (smaller than around 15 bp). Larger insertion/deletions, as well as structural variants, have not been within the scope of this study. It is possible that the genomic region contains such variants, and that they may be directly or indirectly linked to the underlying causative mutation.

The identify of the underlying causative mutation has significance in one other aspect, due to the difference in genetic architecture of SRS resistance in coho versus Atlantic salmon. In Atlantic salmon the trait is under polygenic control (e.g. Marín-Nahuelpi et al. 2024) whereas in coho salmon it is strongly influenced by the QTL on chromosome 21. Thus, one could envision “copying” the resistance mechanism of the coho-QTL onto Atlantic salmon, to increase the resistance levels of Atlantic salmon. Phylogenetic analysis showed that the Hint3 protein encoded by LOC109866665 forms a distinct clade, having no close paralog/ortholog in coho salmon or Atlantic salmon. Thus, the protein is likely to have a unique function not covered by another gene. The lncRNA encoded by LOC109866666 is not present in Atlantic salmon (Figure 10). Therefore, one might need to copy the entire cDNA of one or both genes to confer SRS-resistance to Atlantic salmon, i.e. transgenics would probably be necessary. There would be no guarantee that such an action would lead to increased resistance, as the function of the resistance allele/gene might be context specific. Also, commercial gene editing of animal species is currently not permitted by the legislations of most countries. Nevertheless, gene editing to achieve disease resistance is a possibility which needs to be considered for the future, given the impact that diseases such as SRS have on aquaculture in Chile and worldwide.

In conclusion, we have identified hint3 and a neighbouring long non-coding RNA gene as candidate causal genes, underlying a major QTL for resistance against SRS in coho salmon. The genes were first implicated by positional overlap with the most significant SNPs from a very fine-grained GWAS. Next, both genes were found to be differentially expressed according to genotypes at the said most significant SNPs. One of the two most significant SNPs may be a quantitative trait nucleotide (QTN) because it has a large impact of the Kozak consensus sequence at the *hint3* gene. Our current results do not suffice to prove causality or to say whether the causal gene is *hint3,* the lncRNA, or both. Other hypotheses also cannot be ruled out. Therefore, further research is needed. In terms of selective breeding, the identified SNPs appear sufficient for very precise marker-assisted selection in coho salmon, across populations. In the future, it might be possible to copy the resistance mechanism of the QTL from coho salmon to Atlantic salmon.

## Material & Methods

### Challenge tests

The challenge tests were carried out at the ATC Patagonia experimental facility (Puerto Montt, Chile) or at Centro Aquícola, Fundación Chile (Puerto Montt, Chile). The same test model was employed on all year classes. Smolts (Table 1) was transported to the test facilities, then split into two groups corresponding to shedders and cohabitants. Each group was held in 7 m³ tanks under controlled conditions (temperature 13–14 °C; salinity 30–33 ppt; oxygen 80–110 % saturation; 24 h photoperiod).

Following acclimatisation to sea water (2 weeks), fish were intraperitoneally injected with 0.1 mL of a *P. salmonis* inoculum (strain PM-18856, LF-89-like, provided by ADL Diagnostic Ltda., Puerto Montt, Chile), while cohabitants were exposed by cohabitation. The challenge tests lasted approximately 60 days (see Table 1), with daily monitoring of mortality. Dead fish were weighed, measured, necropsied, and tissue samples (kidney and liver) from 10 % of cohabitant mortalities were preserved in ethanol for confirmatory analysis. For year classes 2019 and 2021, fish were also weighed before the challenge.

At the end of the trial, all surviving fish were euthanized using anaesthetic overdose. Data collected included survival, time-to-death, and biometric measures. Tissue samples were taken from fins. All waste material, including fish remains and inoculum residues, was disposed of according to Chilean biosecurity regulations.

### Genetic analyses

DNA was extracted from fin biopsies using commercially available DNA extraction kits. DNA was genotyped using an Axiom myDesign SNP-genotyping array from Thermofisher (Waltham, MA, USA). The array was custom-made by AquaGen, was named Oki72kv2, and contained 70,571 unique SNPs. The array was a modified version of another Axiom myDesign array named Oki72k, containing 72,015 SNPs. The development of Oki72kv2 from Oki72k entailed 1) removing 10429 SNPs which had displayed suboptimal genotype quality, and 2) adding 8985 SNPs which were in strong to moderate (r^2^ > 0.4 in a dataset derived from 91 whole-genome sequenced coho salmon of diverse ancestry) LD with the estimated genotyped at the SRS QTL (Moen et al. 2022). Oki72k was derived from Oki200k, an Axiom myDesign array containing 200,000 SNPs which had been made on the basis of Illumina sequencing (15x coverage) of a set of 20 diverse coho salmon individuals. Genotyping of Axiom myDesign arrays was according to standard manuals by Thermofisher. SNP-calling from raw data was done using APT Power Tools (Thermofisher) with a culling threshold (for samples) of 0.8 during Dish-QV. In the subsequent genotype-calling process, samples were culled if they had a call rate below 0.97. After SNP-calling, SNPs went through a filtering process where: 1) SNPs were removed entirely of they were not PolyHighResolution, NoMinorHom, or MonoHighResolution in 90% or more of all batches of genotyped fish (using all available coho data). 2) Individual genotypes were culled if their confidence statistic was above 0.01. 3) All genotypes for a SNP were set to missing within a batch of genotyped samples if the SNP was notPolyHighResolution, NoMinorHom, or MonoHighResolution within that batch. 4) Missing genotypes were imputed using Fimpute with standard settings (without use of pedigree information).

Heritabilities were calculated within dataset using the –reml option of GCTA (Yang et al. 2011). Genetic correlations between DPC and survival within dataset was calculated using the –reml-bivar option of GCTA.

The across-datasets GWAS (meta-GWAS) was also done using the –mlma-loco option of GCTA, where dataset was accounted for by including one binary indicator variable as covariate for each dataset.

GWAS and heritability estimations were performed with initial body weight as covariate. This was done only on datasets C19co, C2co, and C21sh, because it was only in these datasets that initial body weight (measure before challenge) was available. The body weigh covariate had a negligible effect on the results. Therefore, the results presented come from analyses done without body weight as covariate.

### WGS population

91 individual Coho salmon, selected to capture the genetic diversity of the AquaGen breeding population, were chosen for whole genome sequencing (WGS). All individuals were sequenced as 150bp PE reads with approximately 10x coverage on an Illumina sequencing machine.

### WGS data preparation

Mapping, variant calling and quality control were performed using the following pipeline / parameters: Illumina reads were mapped to the Coho salmon genome (Okis_V2.1; GCA_002021735.2) using *bwa-mem* (Li and Durbin 2009). Duplicate reads were marked with *samblaster* (v0.1.24) (Faust and Hall, 2014). Genetic variants were identified per sample using GATKs *HaplotypeCaller* (McKenna et al. 2010) applying the default parameters. All samples were subsequently merged into .vcf using GATKs *GenomicsDBImport* / *GenotypeGVCFs*. To avoid detection of false positives the variants were filtered according to the following conditions: i) ‘QD < 2.0’ which filters out variants with a low-quality score by depth of coverage. ii) ‘QUAL < 40.0’ to sort variants with a low-quality score. iii) ‘SOR > 3.0’ removes variants where the number of reads supporting the reference allele is significantly skewed towards one strand (forward or reverse) compared to the other. iv) ‘MQRankSum < −12.5’ removes variants where mapping qualities of REF vs ALT are heavily skewed. v) ‘ReadPosRankSum < −8.0’ removes variants that tend to be near the end of all supporting reads. vi) ‘FS >60.0’ to filter out variants whose reads supporting the reference allele versus the alternative allele are significantly different between the two strands. vii) ‘ReadPosRankSum < −8.0’ to remove variants with a low score for the difference between the mean position of the reference allele and the alternative allele among the reads supporting the variant. All steps were conducted in accordance with the GATK Best Practice document.

Functional annotation of the identified variants was performed using the Ensembl Variant Effect Predictor (VEP). Variants were classified based on their genomic context (e.g., intronic, exonic, intergenic) and predicted impact on protein coding sequences (e.g., synonymous, missense) using the Coho salmon genome annotation (GCF_002021735.2).

Linkage disequilibrium (LD) was estimated as the squared correlation coefficient (r^2^) between the two most significant GWAS SNPs (positions 22,837,421 and 22,842,793) and all other bi-allelic variants within a 5 Mb window, using VCFtools v0.1.16. (Danecek et al. 2011).

### RNAseq

Tissue panel: Samples for a tissue panel were collected from Coho salmon from the AquaGen population, snap frozen in liquid nitrogen and stored at –80°C until final processing (Table 5). Head kidney samples were collected from 19 Atlantic salmon (Salmo salar) belonging to 19 different full-sib families (one fish per family) taken from the yc 2014 challenge test. Ten fish exhibited clinical signs of disease at peak mortality, while the remaining nine appeared healthy (control group). Fish were euthanized by overdose of benzocaine prior to sampling. Immediately after collection, tissue samples were immersed in RNAlater® Stabilization Solution (Thermo Fisher Scientific) and stored at 4°C overnight, followed by long-term storage at −80°C until final processing. Total RNA was extracted individually from each sample using TRIzol® Reagent (Thermo Fisher Scientific) according to the manufacturer’s protocol. Genomic DNA contamination was removed by on-column DNase I digestion using the RNase-Free DNase Set (QIAGEN). RNA integrity was evaluated using the Fragment Analyzer (Advanced Analytical Technologies/BioAnalyzer); all samples exhibited RNA Integrity Numbers (RIN) ≥ 8.0, confirming high-quality RNA suitable for sequencing.

**Table 5:** Overview over the number of biological samples used for ATAC and RNAseq.

|  | RNAseq |  |  |  | ATACseq |  |  |  |
| --- | --- | --- | --- | --- | --- | --- | --- | --- |
|  | Immature Female | Immature Male | Mature Female | Mature Male | Immature Female | Immature Male | Mature Female | Mature Male |
| Brain | 3 | 3 | 3 | 3 | 1 | 1 | 1 | 1 |
| DistalIntestine | 2 | 3 | 3 | 3 |  |  |  |  |
| Gill | 3 | 3 | 3 | 3 | 1 | 1 | 1 | 1 |
| Gonad | 3 | 3 |  |  | 1 | 1 |  |  |
| HeadKidney | 3 | 3 | 3 | 3 |  |  |  |  |
| Liver | 3 | 3 | 3 | 3 | 1 | 1 | 1 | 1 |
| Muscle | 3 | 3 | 3 | 3 | 1 | 1 | 1 | 1 |
| Skin | 2 | 2 | 2 | 3 | 1 | 1 | 1 | 1 |
| Spleen | 3 | 3 | 3 | 3 |  |  |  |  |

RNAseq library preparation as well as sequencing for the tissue panel samples was conducted by NovoGen. RNAseq libraries from head kidney samples were processed in-house using the TruSeq® Stranded mRNA Library Prep Kit (Illumina) following the manufacturer’s instructions. Briefly, mRNA was fragmented, reverse-transcribed into first-strand cDNA using random hexamers and SuperScript II (included in the kit), followed by second-strand cDNA synthesis, end repair, 3′-adenylation, indexed adapter ligation, and PCR enrichment (15 cycles). Library quality and quantity were assessed using the Fragment Analyzer and Qubit® dsDNA HS Assay (Thermo Fisher Scientific). Indexed libraries were pooled equimolar and sequenced on an Illumina MiSeq platform using a 75 bp paired-end configuration (v3 chemistry) at the Laboratory of Biotechnology and Animal Genomics (FAVET-INBIOGEN), Faculty of Veterinary and Animal Sciences, University of Chile, Santiago, Chile.

### RNAseq data processing

RNA sequencing files (.fastq) were processed using a custom pipeline using the following steps: (i) Raw paired-end (PE) reads were quality controlled using *FastQC* (ii) The raw PE reads were subsequently quality trimmed using *cutadapt* v 4.1 (Martin et al.,2011), removing traces of the Illumina 3’ and 5’ adapter, trimming low-quality bases from the end of the reads (parameter -q 25) and only keeping reads with a minimum length (iii) Trimmed PE reads were again quality checked using *FastQC* (iv) Trimmed PE reads were quantified using *salmon* using the Coho transcriptome *. (v)* Trimmed PE reads were aligned against the Coho genome (Okis_V2, GCF_002021735.2) using *STAR in* two-pass mode (Dobin et al. 2013). Further variant calling on the RNAseq data was performed following GATK Best Practices for RNA-seq short variant discovery (https://gatk.broadinstitute.org/hc/en-us/articles/360035531192): Duplicate marking, base quality score recalibration (when applicable), and variant calling were performed using GATK v4.6. Joint genotyping across all 19 successfully sequenced samples was carried out using GenomicsDBImport and GenotypeGVCFs, resulting in a multi-sample VCF.

To investigate whether SNPs in the genomic within or in the vicinity of differentially expressed genes could act as cis-regulatory elements, we performed permutation-based cis-eQTL mapping. We focused on a 200 kb window (±100 kb) centered on the transcription start site (TSS) of the SNP (TSS coordinates extracted from the GTF annotation of the Okis_V2 assembly).

Allele dosages (0/1/2) for each SNP were extracted from the joint-genotyped VCF. Only SNPs passing GATK hard filters and with minor allele frequency (MAF) ≥ 0.05 were retained. For each gene–SNP pair within the 200 kb window, a poisson generalized linear model was fitted using normalized gene counts (from DESeq2) as the response and genotype dosage as the predictor:

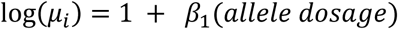

where *β*_1_ represents the per-allele effect on gene expression. We also fit a dominance by including two categories (no resistance allele versus more tha 1 allele). The model is:

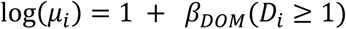

Where *D_i_* indicator equals 1 if individual has at least one alt allele (heterozygous or homozygous), 0 otherwise; *β_DOM_* effect of carrying at least one copy (vs. none). Thus this model assume that Heterozygote (1 copy) and homozygote (2 copies) have the same expected expression

Model fitting was performed using ‘statsmodels’ in Python. Empirical p-values were obtained by permuting genotype labels 100,000 times while keeping expression values fixed. Resulting p-values were adjusted for multiple testing using the Benjamini–Hochberg false discovery rate (FDR) procedure.

## Acknowledgments

The authors would like to thank Dr. Gareth B. Gillard for his excellent support in handling of the ATACseq as well as RNAseq data. The study are also grateful to the LICEResist project and its project leader Prof. Sigbjørn Lien, for making the data available. The study was performed with financial support from CORFO.

